# Topographical differences during motion processing in autistic and dyslexic children

**DOI:** 10.64898/2026.01.30.702841

**Authors:** Etienne Sallard, Anna Esenther, Gaia Scerif, Pawel J. Matusz, Catherine Manning

## Abstract

Atypical motion processing has been reported in multiple developmental conditions, including autism and dyslexia, and taken as support for a general developmental vulnerability in the dorsal stream. Yet by uncovering dynamically unfolding processes, electroencephalography can determine the extent to which motion processing signatures are shared or distinct across different developmental conditions. Here we used pre-registered topographical analyses across the whole high-density electrode array to determine, for the first time, whether autistic and dyslexic children activate the same or distinct brain pathways as typically developing children, when completing two motion processing tasks. Participants were 29 autistic, 44 dyslexic and 57 typically developing children aged 6 to 14 years. Group differences in overall brain response strength were found in both tasks. Topographical activity differed in both autistic and dyslexic children compared to typically developing children (but not between autistic and dyslexic children) in a motion coherence task between 456 - 560 ms. However, group differences in a direction integration task (with no incoherent motion) depended on stimulus difficulty and time window (434 - 500 ms and 538 - 638 ms), with different patterns of divergence from typical development in autism and dyslexia. These results suggest that there are differences in the brain networks used to accomplish motion processing tasks in both autistic and dyslexic children, and demonstrate the utility of a topographical approach for detecting group differences in neural mechanisms, which can be missed by univariate approaches.

**Key points:**

- We use an electrical neuroimaging approach to provide new insights into the neural mechanisms of motion processing in autism and dyslexia
- Autistic and dyslexic children both differed from typically developing children in the strength and topography of their brain responses.
- Motion processing activates atypical brain networks in both autistic and dyslexic children.

## Introduction

Studies of the visual system can give us a ‘window’ into the developing brain (Braddick et al., 2011), as the neural mechanisms underlying the developed visual system are relatively well understood, and we have sensitive methods to probe changes across the lifespan. One interesting observation is that the dorsal stream, which is heavily involved in position and motion processing, develops more slowly than the ventral stream, which is associated primarily with form processing (Braddick, 1993; Braddick et al., 2016). It has also been reported that dorsal stream functions are disproportionately affected relative to ventral stream functions in a range of different developmental conditions (Braddick et al., 2003). These findings have led to the proposal that the dorsal stream is particularly vulnerable to atypical development (*dorsal stream vulnerability*; Braddick et al., 2003).

A common measure used to index dorsal stream functioning is the motion coherence threshold (Newsome & Pare, 1988). The motion coherence task is a global motion task, which requires integrating over space and time, and involves area MT/V5 (Britten et al., 1992). In this task, a proportion of coherently moving ‘signal’ dots are displayed amongst randomly moving ‘noise’ dots, and participants are asked to determine the overall motion direction. The coherence threshold is derived as the proportion of signal dots required to reach a certain level of accuracy in a discrimination or detection task. Motion coherence thresholds are elevated in a range of different developmental conditions (Atkinson et al., 1997; Benassi et al., 2010; Kogan et al., 2004; Van der Hallen et al., 2019), which suggests similar effects across developmental conditions, in accordance with dorsal stream vulnerability.

In this study we focus particularly on autism (a developmental condition characterised by differences in social communication and repetitive and restricted behaviours and interests; World Health Organisation, 2019), and dyslexia (a developmental condition characterised by reading and spelling difficulties; Carroll et al., 2025). Meta-analyses have established elevated motion coherence thresholds in both autistic (Van der Hallen et al., 2019) and dyslexic populations (Benassi et al., 2010) relative to those without developmental conditions (typically developing: TD). Despite this similarity, work using additional tasks and computational modelling has suggested important differences in how autistic and dyslexic people process motion stimuli within the dorsal stream.

First, Pellicano and Gibson (2008) presented a flicker contrast test alongside the motion coherence task, to investigate the functioning of the dorsal pathway at lower, subcortical levels. They found that autistic children had similar flicker contrast thresholds, but elevated motion coherence thresholds relative to TD children, suggesting differences emerging only at higher levels of the dorsal stream. Conversely, dyslexic children had elevated thresholds for both tasks relative to TD children, suggesting differences that originate at lower levels of the dorsal stream. Second, Manning et al., (2015, 2022) presented a direction integration task where the dot directions in each stimulus were drawn from a Gaussian distribution, and participants were asked to discriminate the overall (i.e., mean) motion direction. The important difference between this task and the motion coherence task is that here there are no randomly moving noise dots that need to be filtered out: the optimal strategy is to average all dot directions. Interestingly, in this direction integration task, autistic children were able to discriminate motion direction over a greater range of directional variability (larger standard deviations in dot directions) than TD children (Manning et al., 2015, 2017). In a computational model (the equivalent noise model), this difference was reflected in *increased* global sampling across dots in autistic children (cf. accounts of reduced global processing in autism; Happé & Frith, 2006). However, dyslexic children did not show the same pattern of results. Instead, dyslexic children showed no evidence of differences in their sampling relative to TD children, but some evidence of higher internal noise estimates, reflecting more imprecise estimates of each dot’s direction (Manning, Hulks, et al., 2022).

Finally, in another set of studies, autistic, dyslexic and TD children discriminated the direction of motion coherence and direction integration stimuli presented at two difficulty levels (Manning, Hassall, et al., 2022b, 2022a). Children’s accuracy and response times were modelled with the diffusion model (Evans & Wagenmakers, 2020; Stone, 1960) to break down performance into distinct processing stages. Here, it was found that dyslexic children showed a slower rate of accumulation of motion evidence towards a decision bound than TD children in both tasks (Manning, Hassall, et al., 2022b). Meanwhile, autistic children showed no differences from TD children in diffusion model parameters (Manning, Hassall, et al., 2022a).

Given these differences between autistic and dyslexic children in behavioural indices and latent parameters, we might expect the neural mechanisms underlying motion processing to differ between autistic and dyslexic children. While few studies directly compare neural responses to motion in autistic and dyslexic, Toffoli et al. (2021) compared evoked responses to the onset of motion coherence and direction integration stimuli in a subset of the TD, autistic and dyslexic children analysed in Manning, Hassall, et al., (2022b, 2022a). Using a data-driven technique, Toffoli et al. identified two sets of electrode weights (or EEG components) that maximised trial-by-trial reliability: 1) an early component that was maximal over occipital electrodes and had a negative peak at ∼160 ms (resembling the motion-specific N2, Niedeggen & Wist, 1998, 1999), and 2) a later, more sustained component which was maximal over centro-parietal electrodes (resembling the centro-parietal positivity, Kelly & O’Connell, 2013; O’Connell et al., 2012). Component waveforms were computed for each participant, by averaging the activity across all electrodes in a weighted fashion, according to the component’s electrode weights. Mass univariate statistics (Groppe et al., 2011) did not uncover group differences in component waveforms at early timepoints including the N2-like peak. However, both autistic and dyslexic children showed significantly increased amplitudes relative to TD children in the occipital component waveform from around ∼430 ms after stimulus onset, specifically for the motion coherence task and not the direction integration task. This effect was speculatively linked to difficulties with segregating signal-from-noise in autistic and dyslexic children. Exploratory analyses suggested some differences between autistic and dyslexic children in response-locked activity in the centroparietal component.

This response-locked EEG activity was more comprehensively dealt with in Manning, Hassall, et al. (2022a, 2022b), who used linear deconvolution to unmix stimulus and response-locked activity. They found that there was a shallower build-up of amplitude in the centro-parietal component preceding the response for dyslexic children compared to TD children, for both motion coherence and direction integration tasks (Manning, Hassall, et al., 2022b). Moreover, this build-up was correlated with reduced evidence accumulation in the diffusion model. Meanwhile, autistic children did not clearly differ in the rate of build-up of amplitude in the centro-parietal component compared to TD children (Manning, Hassall, et al., 2022a).

In summary, previous analyses show both similar and distinct features in autistic and dyslexic children’s component waveforms during motion processing tasks. However, by averaging across electrodes to form component waveforms and conducting univariate analyses, we lose information about whether autistic and dyslexic children are activating the same brain pathways as TD children, but to a greater or lesser extent, or whether they are relying on different brain pathways. Recent advances in multivariate electrical neuroimaging (Habermann et al., 2018) allow us to discriminate between these possibilities by interrogating group differences across the whole scalp electric field. Specifically, electrical neuroimaging yields two robust, reference-independent measures directly interpretable in terms of neurophysiological mechanisms: brain response strength (“gain control”, through Global Field Power; GFP) and the recruited brain region configurations (through topographical analyses of variance, TANOVA; Koenig et al. 2011; Lehmann & Skrandies, 1980; Michel & Koenig, 2018; Matusz et al. 2019). In this study, we therefore reanalysed the data from Toffoli et al. (2021) using electrical neuroimaging to provide new insights into the neural mechanisms involved in motion processing in autistic and dyslexic children.

### Research questions and hypotheses

We had two pre-registered research questions and hypotheses (https://osf.io/fy938/overview):

*1. Do autistic and dyslexic children differ in response strength (GFP) compared to TD during motion coherence and direction integration tasks?*

We hypothesised a main effect of group in GFP, indicating group differences in the strength of the response. Previous reports of increased V5/MT activation in autism (Peiker et al., 2015; Brieber et al., 2010; Nyström et al., 2021; Schallmo et al., 2020) and reduced activation in dyslexia (Olulade, Napoliello, & Eden, 2013; Demb, Boynton, & Heeger, 1997) led us to expect that GFP might be increased in autism, but reduced in dyslexia in the motion coherence task. Previous research has not examined V5/MT activation during motion integration, so there was less evidence to suggest a GFP difference in this task.

*2. Do autistic and dyslexic children demonstrate the use of different sources of activation (measured by TANOVA) than TD children during motion coherence and direction integration tasks?*

We hypothesised a main effect of group in the TANOVA analysis, indicating a qualitative difference in the underlying sources of activation between groups. This hypothesis is based on preliminary evidence from previous studies of possible topographical differences. Infants at increased likelihood of autism show more lateralised processing during motion coherence processing than those at low likelihood of autism (Nyström et al., 2020). Moreover, topographical differences have been reported in response to phonological stimuli in dyslexia (Van Leeuwen et al., 2006; Spironelli, Penolazzi, & Angrilli, 2008, Vourkas et al., 2011). Regarding visual motion processing, Dushanova and Tsokov (2021) suggested that visual motion training can improve reading outcomes and alter the network topologies of dyslexic children to become more like non-dyslexic controls. Although no studies have yet conducted a topographical comparison of autistic, dyslexic and typically developing children during motion coherence and direction integration tasks, we hypothesised that topographical differences might emerge based on these previous studies.

## Materials and Methods

### Participants

Participants were TD children without any diagnosed developmental conditions (n = 57), diagnosed autistic children (n = 29) and children with a dyslexia diagnosis (n = 44, including 1 in the process of obtaining a diagnosis) aged 6 to 14 years (Table 1). Full details of recruitment and inclusion criteria are included in Toffoli et al. (2021). Briefly, all children had normal or corrected-to-normal visual acuity, and verbal and performance IQ over 70, as measured by the Wechsler Abbreviated Scales of Intelligence, 2^nd^ edition (WASI-2; Wechsler, 2011). Children with both autism and dyslexia diagnoses were excluded from the dataset. Autistic children exceeded cut-off criteria for autism on one or both of the Social Communication Questionnaire (SCQ; Rutter et al., 2003) and Autism Diagnostic Observation Schedule, Second Edition (ADOS-2; Lord et al., 2012). Dyslexic children had reading and spelling composite scores of 89 or below (formed from standard scores from the Wechsler Individual Achievement Test, 3^rd^ edition [WIAT-III, Wechsler, 2017] and Test of Word Reading Efficiency, 2^nd^ edition [TOWRE-2, Torgesen et al., 2012] Phonemic Decoding Efficiency). TD children and dyslexic children had SCQ scores under the cut-off (15), and TD children and autistic children had reading and spelling composite sores above 89. As shown in Table 1, the dyslexic children had a lower mean IQ than the other groups, but we decided not to control for IQ in our analyses following Dennis et al. (2009), as it might be related to both dyslexia and motion processing. Data from one child from each group are missing from the direction integration task following technical issues or the child’s request to complete that task without EEG.

**Table 1.** Participant demographics

| Demographic | Typically developing | Autism | Dyslexia |
| --- | --- | --- | --- |

| variable |  |  |  |
| --- | --- | --- | --- |
| Sex (M:F) | 32:25 | 22:7 | 19:25 |
| Age | 10.50 (2.22) 6.55 –<br>14.98 | 11.04 (2.57) 6.54 -<br>14.94 | 11.02 (1.79) 8.26 –<br>14.53 |
| Verbal IQ | 111.49 (9.19) 95 –<br>132 | 109.76 (12.59) 85 –<br>137 | 100.23 (9.75) 82 –<br>118 |
| Performance IQ | 112.46 (13.37) 81 –<br>145 | 109.55 (14.59) 78 –<br>136 | 101.2 (15.22) 72 –<br>141 |
| Full-scale IQ | 113.53 (10.21) 89-135 | 110.9 (13.21)<br>84-133 | 100.64 (12.18)<br>79-132 |
| SCQ | 2.56 (2.75) 0 – 12 | 19.69 (7.52) 4 – 32 | 4.91 (3.74) 0 – 14 |
| TOWRE-2 PDE | 106.95 (10.91) 80 –<br>135 | 107.76 (11.83) 86 –<br>132 | 78.57 (6.97) 65 – 99 |
| WIAT-III spelling | 113.3 (16.33) 84 –<br>153 | 106.86 (18.9) 68 –<br>152 | 79.73 (8.19) 59 – 99 |
| Reading and<br>spelling composite | 110.12 (12.55) 89.5<br>– 140.5 | 107.31 (12.88) 89.5<br>– 142 | 79.15 (6.12) 63 –<br>88.5 |
| ADOS-2 Total | - | 12.17 (5.37) 4 – 27 | - |

Notes. SCQ = Social Communication Questionnaire; TOWRE-2 = Test of Word Reading Efficiency, 2<sup>nd</sup> edition; PDE = Phonemic Decoding Efficiency; WIAT-III = Wechsler Individual Achievement Test, 3<sup>rd</sup> Edition. ADOS-2 = Autism Diagnostic Observation Schedule, 2<sup>nd</sup> edition.

The study was approved by the Central University Research Ethics Committee at University of Oxford. Parents provided written informed consent and children provided written assent.

## Stimuli and apparatus

Stimuli were 100 white dots (0.19° diameter) plotted in random positions within a 10° x 10° square aperture on a black background. The dots moved at 6°/s with a limited lifetime of 400 ms. A central red fixation square (0.24° x 0.24°) was displayed. Stimuli were displayed on a Dell Precision M3800 laptop (2048 x 1152 pixels, 60 Hz) using MATLAB and the Psychophysics Toolbox (Brainard, 1997; Pelli, 1997). Responses were recorded with a Cedrus RB-540 response box. EEG data were collected with 128-channel Hydrocel Geodesic Sensor Nets and Net Amps 300, using NetStation 4.5 software (Electrical Geodesics Inc., OR, USA).

## Task procedure

Participants sat at a viewing distance of 80 cm in a dimly-lit, electrically shielded room and completed motion coherence and direction integration tasks in a counterbalanced order. They were told that ‘fireflies’ (dots) were escaping from their boxes, and that they had to help the zookeeper determine which direction they were escaping in, as quickly and accurately as possible.

Figure 1 shows the structure of each trial, including a fixation period, random motion period, stimulus period, and offset period. The random motion period ensured the separation of evoked responses to directional motion from pattern- and motion-onset evoked potentials. An auditory tone marked the start of the stimulus period. In the stimulus period of the motion coherence task, either 75% (easy) or 30% (difficult) of dots moved coherently leftwards or rightwards, while the rest continued to move randomly. In the stimulus period of the direction integration task, the dot directions were taken from a Gaussian distribution with a mean leftward or rightward direction and a standard deviation of 30° (easy) or 70° (difficult). The fixation, random motion and offset periods had random durations within a fixed range. The stimulus period was displayed until the participant’s response or 2500 ms. Directional motion continued in the offset period to separate motion offset from participant response. Participants were given feedback if they did not respond within 2500 ms. For each task, 144 trials were presented (72 per difficulty level, with equal rightward and leftward trials), with 8 randomly interleaved catch trials (with 100% coherent / 0° SD motion), split across 4 blocks. Points were given at the end of each block, reflecting overall response time and accuracy. Before these blocks, a practice block was presented for each task, where task difficulty was gradually increased and a criterion of 4 correct consecutive responses was required (see Toffoli et al., 2021). See https://osf.io/wmtpx/ for experimental code.

**Figure 1.**
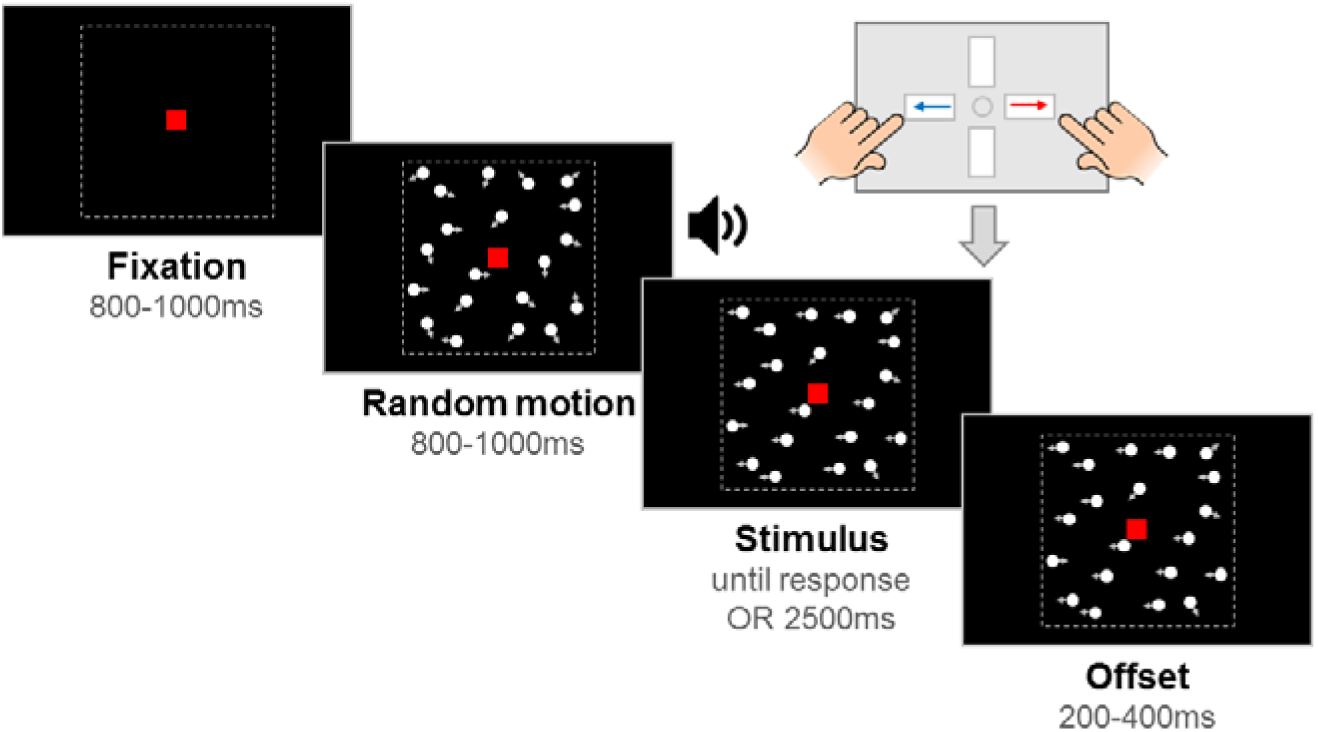
Schematic representation of trial structure. A *fixation* period was followed by a *random motion* period with dots moving in random directions. An auditory tone marked the start of the *stimulus* period introducing directional motion (n.b. the stimulus represented here is from the motion coherence task). After the participant reported the direction with a response box, or the maximum stimulus duration (2500 ms) was reached, the directional motion continued during the *offset* period. Arrows (showing movement) and dotted lines (showing the stimulus aperture) are for illustrative purposes. Figure reproduced from https://osf.io/wmtpx/ under a CC-BY4.0 license.

## EEG procedure

Before the tasks, the EEG net was fitted and electrode impedances were checked. Adjustments were made to keep impedances below 50 kΩ. EEG was sampled at a rate of 500 Hz with a vertex reference electrode. After the first task, impedances were re-checked and adjusted. EEG data quality was monitored throughout the tasks, and adjustments made where possible at the end of blocks.

## General procedure

The EEG set-up and tasks lasted approximately 1 hour. Additionally, children completed a Snellen acuity test, the TOWRE-2 (Torgesen et al., 2012), the WIAT-III spelling subtest (Wechsler, 2017), and the WASI-2 (Wechsler, 2011). Autistic children also completed the ADOS-2 (Lord et al., 2012). A researcher carefully monitored children during all tasks and provided general encouragement and task reminders. Breaks were given as needed.

## EEG pre-processing

We used preprocessed data from Toffoli et al. (2021). Briefly, EEG data were bandpass filtered between .3 and 40 Hz, epoched from the fixation period onset to the end of the offset period and median-corrected for DC offsets. Bad electrodes were identified across each task (≥15% samples above the 97.5th absolute amplitude percentile for each participant) and replaced with the average of the nearest neighbouring electrodes. Horizontal and vertical electrooculogram (EOG) were linearly regressed out from each channel. Channels were removed for a trial if they had ≥15% of samples above the 97.5th absolute amplitude for each participant. Samples that were four or more standard deviations away from the mean were replaced with missing values. EEG data were removed from trials where ≥15% of channels were removed and we removed the data from three electrodes for two participants which had no signal (flat activity). Next, we extracted average event-related potentials for each channel and participant across trials, locked to the stimulus onset from 0 to 800 ms and baselined to the average of the last 100 ms of the random motion period. Finally (and additionally to Toffoli et al., 2021), we interpolated any missing data with the average of neighbouring electrodes, and saved the participant averaged waveforms in .ascii format.

## EEG analysis

Differences in the ERPs across group and task difficulty were analysed using a multi-step analysis approach, known as electrical neuroimaging, which involves global measures of the electric field at the scalp. All analyses were done in Ragu, which in its recent version enables both GFP and TANOVA analyses (for overall info on the software, especially for the TANOVA in earlier versions, see Koenig et al. 2011). The electrical neuroimaging methods have been described in extensive detail previously (e.g., Lehmann & Skrandies, 1980; Michel & Koenig, 2018; Matusz et al. 2019). Our analysis was preregistered (https://osf.io/fy938) following a preliminary run of the analysis steps with a subset of 3 participants from each group, from the motion coherence task, to ensure viability. We imported the data into Ragu using a developmental montage, and used the automatic outlier detection in Ragu to remove outliers based on the Mahalanobis distance, and the data were L2 normalized (Habermann et al., 2018).

Our pre-registered analysis dealt with the data from each task separately. First we investigated whether the factors of group and difficulty modulated the strength of ERP responses within statistically indistinguishable brain networks. For this purpose, we measured the Global Field Power (GFP), which equals the root mean square, or standard deviation, across the average-referenced electrode values at a given moment (as described in Lehmann & Skrandies, 1980). The GFP waveform is a moment-to-moment measure of standard deviation of potential (μV) across the whole montage. However, GFP does not provide insight into how the potentials are distributed across the scalp. Differences in GFP between two conditions “without” concomitant statistically reliable differences in the scalp topography (as measured with Global Dissimilarity, DISS, see below) stem from a change in the gain within statistically indistinguishable generators between two conditions. We remind the reader that GFP and DISS are reference independent.

The GFP waveform can be assessed statistically just like any other ERP waveform. We evaluated the significance of each of the two factors (difficulty and group), and the difficulty by group interaction with three statistics, namely the count of sub-threshold p-values (Global Count Statistics: GCS), the duration of periods of contiguous sub-threshold p-values (Global Duration Statistics: GDS), and Fisher’s combined probability (Area Under the Curve: AUC). Each of these three statistics accounts for the effects of multiple testing by comparing the observed GFP to a distribution of GFPs calculated through 1000 randomization runs. The first two are straightforward. The GCS indicates whether a larger number of timepoints yield a significant p-value than would be expected by chance and the GDS indicates whether contiguous periods of significant p-values are longer than would be expected by chance. Calculating the third statistic, AUC, involves summing the negative natural logs of each p-value, resulting in larger values of the statistic for smaller p-values. Hence, the three statistics give information about the number of significant p-values, the duration of time periods with significant p-values, and the overall distribution of p-values, respectively (Habermann et al., 2018; Fisher, 1925). We used a significance threshold of *p*=0.05.

Next, we conducted a topographic consistency test to confirm that the grand-mean GFP in each condition is higher than it would be by chance for most of the time period. Then, we tested whether the factors modulated the ERPs through modulations in scalp-level field topography and so in the configurations of brain sources that the groups recruited between the two levels of task difficulty across two tasks. Differences between two electric fields (independent of the strength of the two) are indexed by global dissimilarity (DISS). DISS is the root mean square of the squared differences between the potentials measured at each electrode (vs. the average reference), where each is first scaled to unitary strength by dividing it by the instantaneous GFP (Lehmann & Skrandies, 1980). DISS is directly related to the spatial Pearson’s product-moment correlation coefficient between the potentials of the two compared voltage scalp-level topographies. Thus, if two ERP topographies are perfectly inverted, the spatial correlation coefficient value would be −1 at a given moment (i.e., DISS value of 2), and this relationship is expressed by spatial correlation being equal to ((1 − DISS2) / 2). If two ERPs differ in topography but their strength is similar, this directly indicates that the two maps were generated by a different configuration of the brain regions recruited for response. Display and analysis of DISS across time allows for defining periods of stable patterns of ERP activity and the changes in this activity. GFP and DISS are de facto inversely related; when GFP is high, ERP topographic activity is usually stable (i.e., DISS is low), but it changes when GFP is low. DISS across time shows a characteristic pattern. The topographic activity remains stable for tens to hundreds of milliseconds and then changes suddenly to a new configuration, lasting again tens to hundreds of milliseconds. These sequentially organized and highly reproducible configurations are well- known to represent successive steps along the information processing pathway from perception to action (“functional microstates”; Lehmann et al., 1987; Brandeis, Lehmann, Michel, & Mingrone, 1995; Michel & Koenig, 2018; for similar implementations, in adults, e.g., Tivadar et al. 2018, Retsa et al. 2018, Turoman et al. 2021a; in developmental populations: Matusz et al. 2018; Maitre et al. 2017; Turoman et al. 2021b, 2021c).

Following this, we analysed the topographic modulations of the observed ERPs by carrying out two 2 (difficulty level) x 3 (group) mixed topographical analyses of variance (TANOVAs), one for each task. We controlled for multiple testing with the same three statistics as in the GFP analysis – count (i.e., GCS), duration (i.e., GDS), and AUC, with a significance level of p = .05. Any time periods identified as significant by the AUC statistic were further explored with t-maps, which show the difference in average topographies between two conditions. If the TANOVA results identified a main effect of group during a particular time period, for example, we would test the significance of the maps of differences between groups in that time period. In our pre-registration, we planned to next conduct a microstate analysis with the atomize and agglomerate hierarchical clustering model. However, with the change of team members including the addition of those with expertise in topographical approaches, we made the collective decision not to run this segmentation analysis, because the results would be difficult to interpret with such a complicated design, and because the pre-registered research questions do not require this segmentation step.

## Results

### Non-pre-registered analyses: Behavioural data

We had no pre-registered hypotheses relating to behavioural data in this study. However, to help contextualise the pre-registered EEG analyses, Figure 2 presents participants’ accuracy, median response times, and inverse efficiency scores (IES; calculated as mean response time for correct trials divided by the proportion correct; Bruyer & Brysbaert, 2011) for each difficulty level. Analyses comparing accuracy and response time across group and difficulty level were reported in Toffoli et al. (2021). Briefly, there was no significant effect of group, nor interactions between group and difficulty level, on accuracy in the motion coherence task. However, in the direction integration task, there was a significant interaction between group and difficulty level on accuracy, where the groups differed in accuracy in the difficult condition only. Planned contrasts were not significant, but autistic children tended to be more accurate, and dyslexic children less accurate overall than TD children. In the motion coherence task, there were significant group differences in response time, but no interactions with difficulty level and group. While the planned contrasts were not significant, autistic children were generally faster and dyslexic children generally slower than TD children. In the direction integration task, there was no significant effect of group or interaction between group and difficulty level on response times.

**Figure 2.**
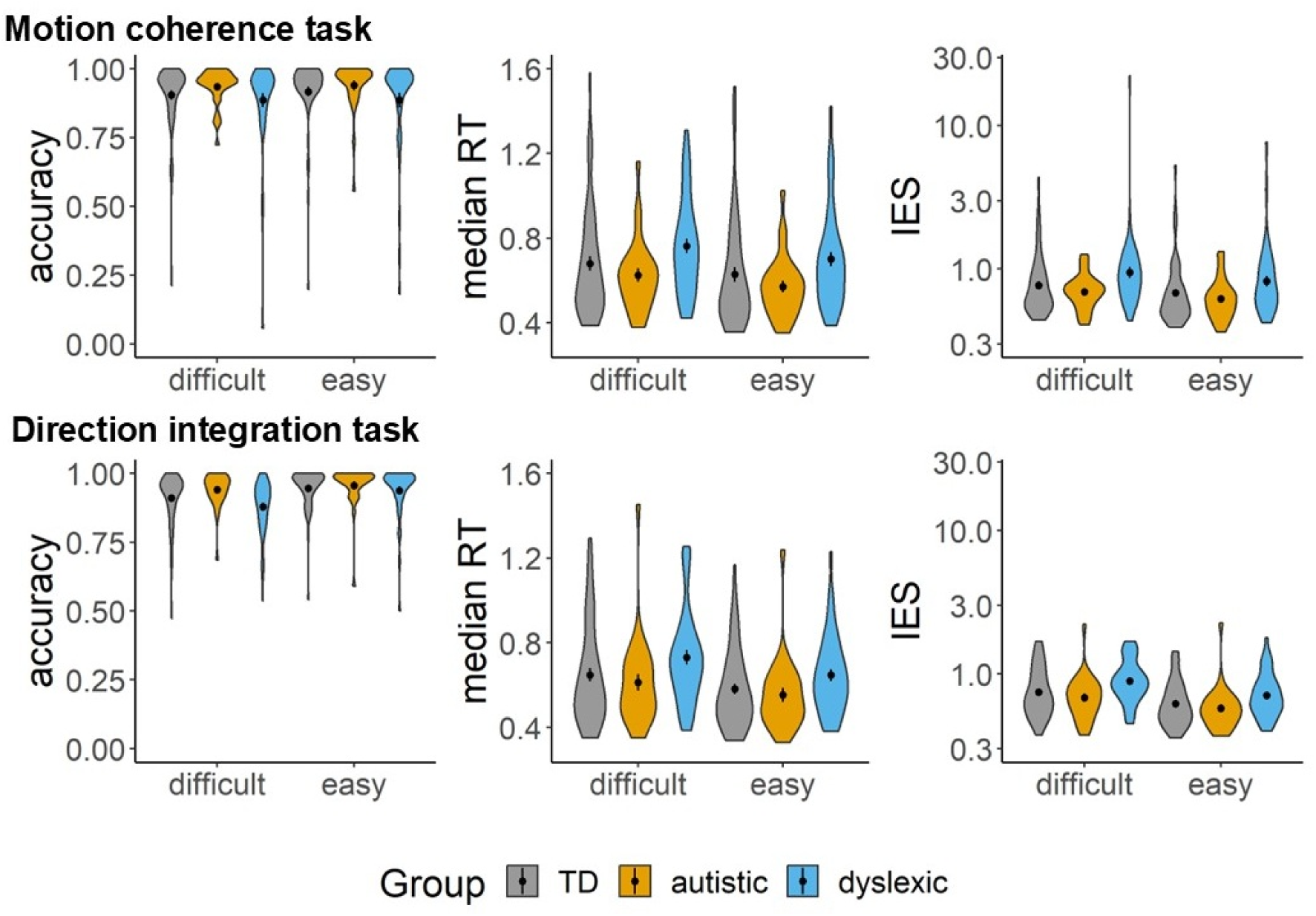
Distributions of accuracy, median response time (RT) for correct trials, and inverse efficiency scores (IES) in motion coherence (top row) and direction integration (bottom row) tasks. The coloured ‘violins’ reflect the kernel probability density of TD (grey), autistic (orange) and dyslexic (blue) participants. The group mean is shown with a black dot ± 1 SEM. Note that IES is presented on a logarithmically spaced axis.

We also explored group differences in log inverse efficiency scores, as a measure that integrates speed-accuracy tradeoffs. In the motion coherence task, there were significant effects of difficulty, *F*(1,127) = 73.77, *p* < .001, ηp2 = .37 (with higher inverse efficiency scores in the difficulty condition), and group, *F(*2,127) = 3.86, *p* = .02, ηp2 = .06. Dyslexic children were less efficient (higher log IES) than both typically developing (*p* = .04) and autistic (*p* = .01) children, while autistic children were not significantly different from typically developing children (*p* = .37). There was no significant interaction between difficulty level and group, *F*(2,127) = .28, *p* = .76, ηp2 < .01. A similar pattern was found in the direction integration task. Log IES was again significantly higher in the difficult condition than the easy condition *F*(1,124) = 339.78, *p* < .001, ηp2 = .73 and the main effect of group was significant, *F*(2,124) = 4.37, *p* = .02, ηp2 = .07. Dyslexic children again had higher log IES than both typically developing (p = .03) and autistic (p = .01) children, with no significant differences between autistic and typically developing children (p = .32). Again, there was no significant interaction between group and difficulty level, *F*(2,124) = 2.26, *p* = .11, ηp2 = .04.

### Pre-registered analyses: Coherence task

For the coherence task, the overall strength at the scalp level analysed with the GFP test showed a main effect of difficulty (GCS: *p* = 0.002; AUC: *p* = 0.001) from 294 to 626ms (GDS= 78ms) and a main effect of group from 628 to 798ms (GDS= 116ms; see Figure 3). Post-hoc comparisons (unpaired t-test with FDR correction; *p-*values averaged over the main effect period of 628-798ms) revealed a significant difference between TD and dyslexic groups (*p* = 0.002) but not between TD and autistic (*p* = 0.14), nor between autistic and dyslexic groups (*p* = 0.22). No interactions were identified.

**Figure 3.**
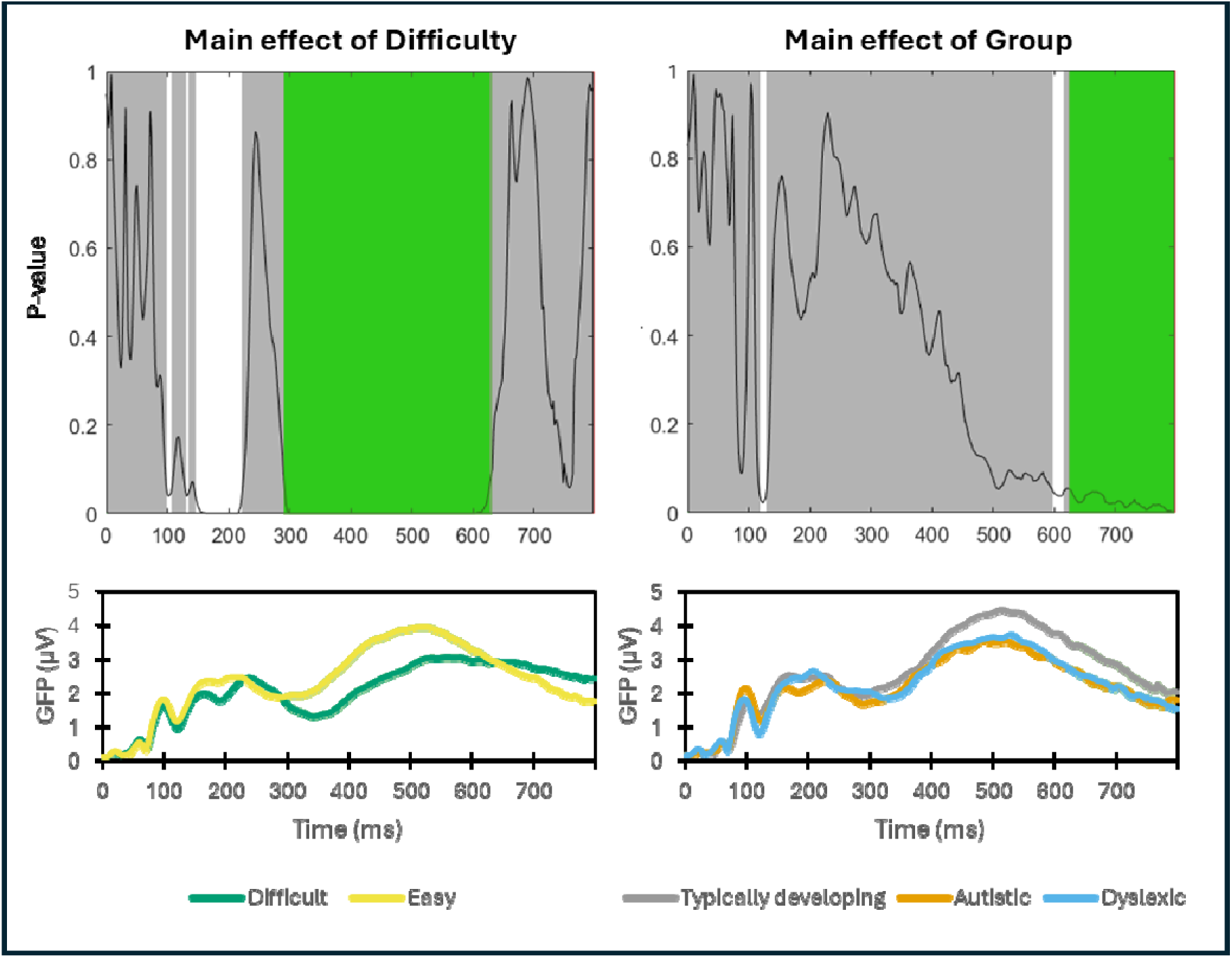
GFP results of the Coherence task analysis. P-value plots (top) are presented for the main effect of difficulty and the main effect of group in whole-epoch analyses of the GFP. The black line represents the p-value, white areas indicate significant time points, and green segments mark intervals where the effect meets global duration criteria (i.e., GCS). The line graph (bottom) represents averaged GFP with the time window of each main effect (main effect of difficulty: 294-626ms; main effect of group: 628-798ms).

The TCT analysis revealed consistent neural activation across participants in each group and for each difficulty level across the entire epoch (Supplementary Figure 1), which indicated that the assumptions were met for running the TANOVA. Finally, the TANOVA analysis showed a main effect of difficulty (GCS: *p* = 0.001; AUC: *p* = 0.001) from 118 to 798ms (GDS= 68ms) and a main effect of group from 456 to 560ms (GDS: 102ms; see Figure 4). Post-hoc comparison (t-maps) for the main effect of group (between 456 - 560ms) showed significant differences between TD and autistic (*p* = 0.008) and TD and dyslexic children (*p* = 0.008) but not between autistic and dyslexic children (*p* = 0.61). No interaction was evidenced.

**Figure 4.**
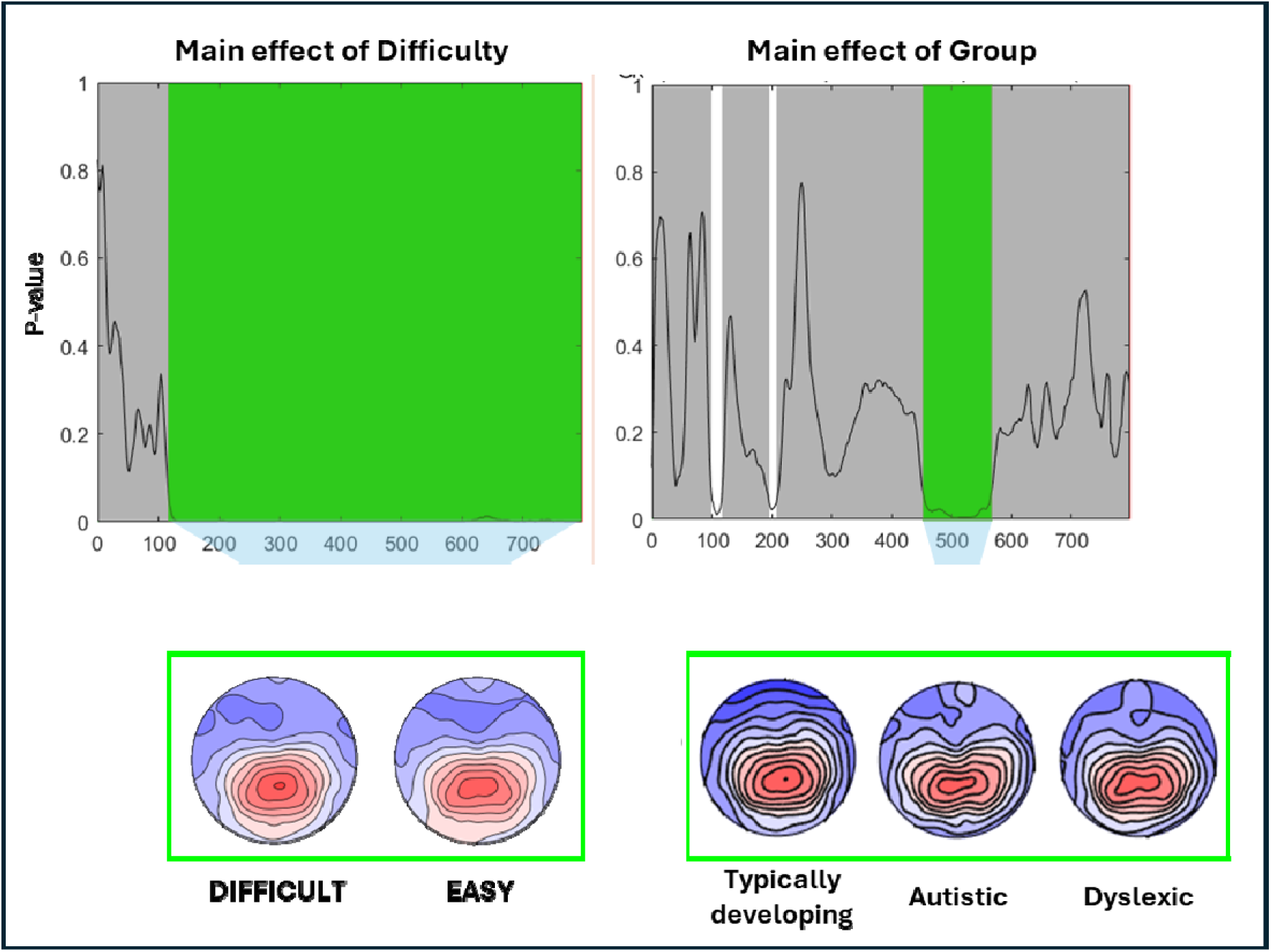
TANOVA results of the Coherence task analysis. P-value plots (top) are presented for the main effect of difficulty and the main effect of group in whole-epoch analyses of the TANOVA. The black line represents the p-value, white areas indicate significant time points, and green segments mark intervals where the effect meets global duration criteria (i.e., GCS). Topographical plots (bottom) within the time window of each main effect (main effect of difficulty : 118-798ms; main effect of group: 456-560ms).

### Pre-registered analyses: Integration task

For the integration task analysis, the GFP analysis showed a main effect of group (GCS: p= 0.078; AUC: p= 0.11) from 556 to 698ms (GDS= 124ms). Post-hoc comparisons (unpaired t-test with FDR correction; p-values averaged over the main effect period of 556-698ms) revealed a significant difference between TD and autistic groups (*p* = 0.02), and between TD and dyslexic groups (*p* = 0.01), but not between autistic and dyslexic groups (*p* = 0.86). and a main effect of difficulty (GCS: *p*= 0.002; AUC: *p* = 0.001) at two periods of time (periods of interest: POI1: 114-224ms and POI2: 330-638ms; GDS= 76ms; see Figure 5). No interaction was evidenced.

**Figure 5.**
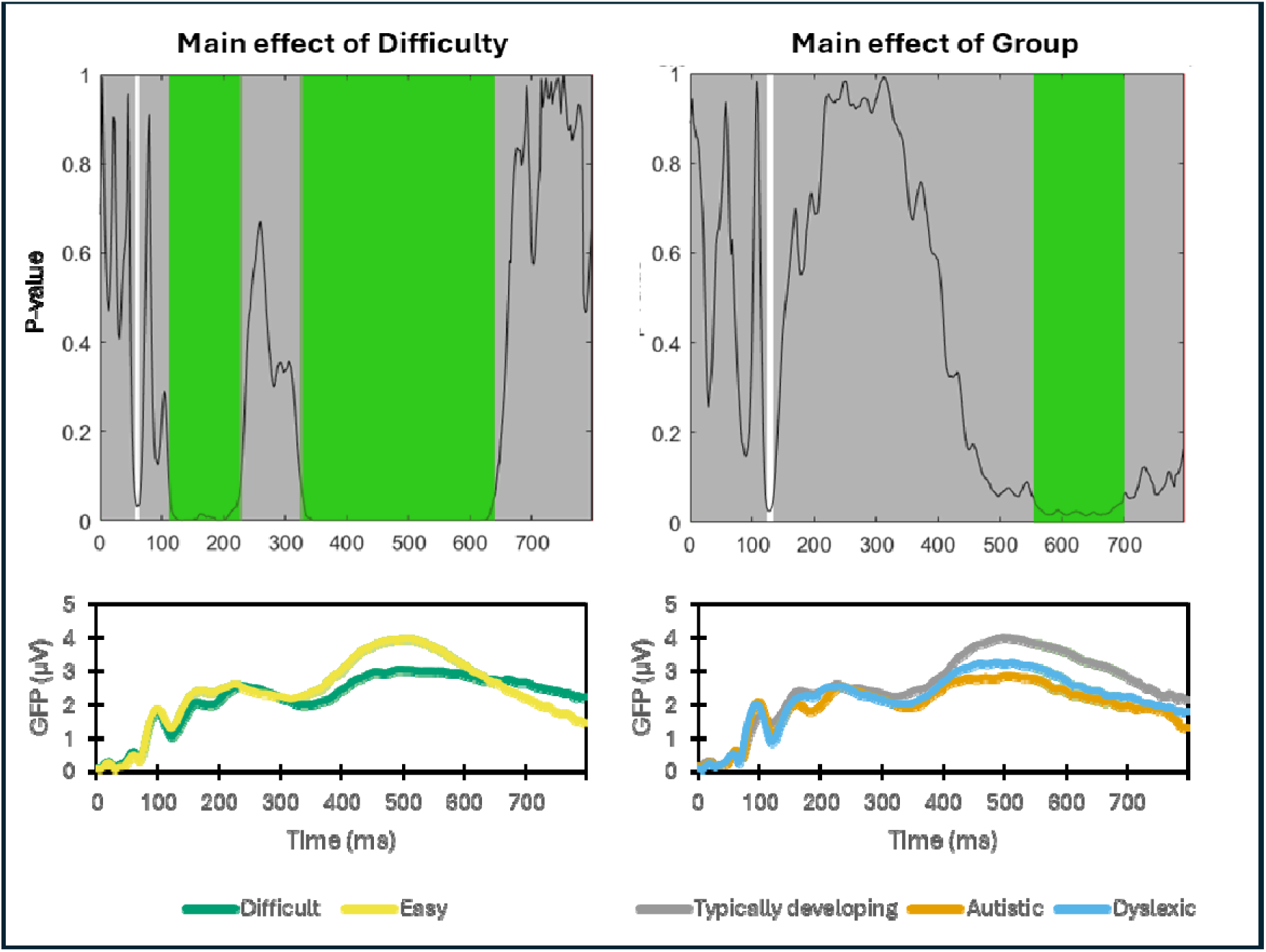
GFP results of the integration task analysis. P-value plots (top) are presented for the main effect of difficulty and the main effect of group in whole-epoch analyses of the GFP. The black line represents the p-value, white areas indicate significant time points, and green segments mark intervals where the effect meets global duration criteria (i.e., GCS). Line graphs represent the averaged GFP (bottom) with the time window of each main effect (main effect of difficulty: POI1: 114-224ms and POI2: 330-638ms; and main effect of group: 556 to 698ms).

TCT revealed consistent neural activation across participants in each group for each difficulty level across the entire epoch (Supplementary Figure 2). Next, the TANOVA analysis showed a main effect of difficulty (GCS: *p* = 0.001; AUC: *p* = 0.001) from 110 to 798ms (GDS= 60ms) and an interaction of difficulty by group (GCS: *p* = 0.03; AUC: *p* = 0.03) at two periods of time (POI 1: 434-500ms and POI 2: 538-638ms; GDS= 60ms; Figure 6). For the POI 1, in the difficult condition, post-hoc comparison showed differences between TD and autistic children (*p* = 0.005) and TD and dyslexic children (p = 0.03) but not between autistic and dyslexic children (*p* = 0.77). For POI 1 in the easy condition, there were differences between TD and dyslexic children (*p* = 0.04) but not between TD and autistic children (*p* = 0.12) nor between autistic and dyslexic children (*p* = 0.18).

**Figure 6.**
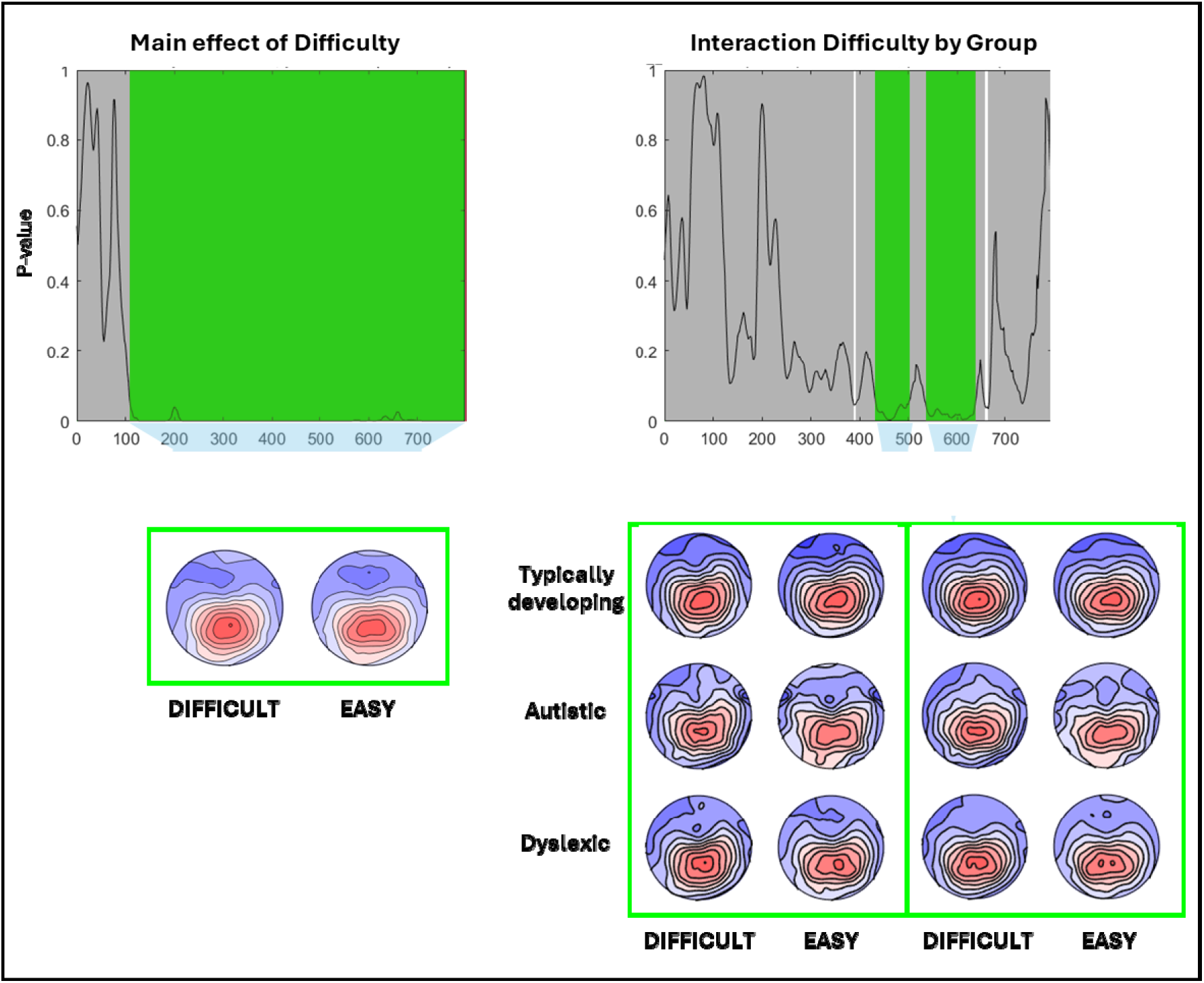
TANOVA results of the integration task analysis. P-value plots (top) are presented for the main effect of difficulty and the main effect of group in whole-epoch analyses of the TANOVA. The black line represents the p-value, white areas indicate significant time points, and green segments mark intervals where the effect meets global duration criteria (i.e., GCS). Topographic maps (bottom) within the time window for the main effect of difficulty (110-798ms) and the interaction of difficulty by group (POI 1: 434-500ms and POI 2: 538-638ms).

For the POI 2 in the difficult condition, post-hoc comparison showed significant differences between TD and autistic children (*p* = 0.001) and a marginal difference between TD and dyslexic children (*p* = 0.054). No significant difference was observed between autistic and dyslexic children (*p* = 0.89). For the easy condition, the differences were found again only between TD and autistic children (p = 0.04) but not between TD and dyslexic children (p = 0.41) nor between autistic and dyslexic children (p = 0.16). No main effect of group was evidenced.

### Non-pre-registered analyses: Task differences

We supplemented our pre-registered analyses on each task with additional 3-way analyses that included task as a factor, to investigate whether different brain networks are recruited by children across the two tasks. We found significant task differences in GFP from ∼150 ms to 500 ms, and task differences in the TANOVA from ∼110 ms to the end of the epoch. Full results are presented in Supplementary Figures 3 and 4.

## Discussion

This study aimed to identify and characterise differences in neural mechanisms between typically developing, autistic and dyslexic children during two motion processing tasks. In our multivariate “electrical neuroimaging” analyses we employed robust, reference-independent measures of brain response strength (a “gain control” mechanism) and topography (measuring differences in the recruited brain region configurations). With this analytical approach, we were able to extend previous insights from work that has compared component average waveforms across the three groups (Toffoli et al., 2021). As hypothesised, we found group differences both in overall brain response strength and topographies, although brain response strength was more similar between autistic and dyslexic children than we had predicted.

The first task we investigated was a coherence task, where the stimulus included either a low (difficult) or high (easy) proportion of coherently moving dots moving among randomly moving dots. In this task, we found no interactions between difficulty and group, but evidence for both main effects, across both GFP and the TANOVA (that is based on the DISS). Specifically, task difficulty modulated the engaged brain networks from ∼120ms post-stimulus onset to the end of the epoch (800 ms post-stimulus), and the overall brain responses became stronger for the easy condition compared to the difficult condition from ∼300ms until ∼630ms post-stimulus. These results suggest that task difficulty elicited the recruitment of statistically distinguishable brain networks, already at the early perceptual-discriminative stages, followed soon after by, and potentially eliciting, the modulations in “gain”-like brain mechanisms across difficulty levels (see, e.g., Matusz et al. 2019, for in-depth discussion). The influence of task difficulty on both mechanisms extends to late cognitive (e.g., decisional) processes. Group differences in the coherence task emerged relatively late, in both strength- and network-based mechanisms. The groups started recruiting distinct brain regions from ∼450ms to 560ms post-stimulus, after which, from ∼630ms post-stimulus to the end of the epoch, the brain responses of dyslexic children were significantly attenuated compared to those of TD children. Brain responses were also slightly attenuated, but not significantly, in autistic children across this period. In this task, the responses of autistic and dyslexic children were non-significantly different across both strength-based and topographic results. Interestingly, Toffoli et al. previously reported group differences between ∼430ms and ∼570 ms post-stimulus, with autistic and dyslexic children both showing *increased* amplitudes relative to typically developing children in a data-driven component with maximal activity over occipital electrodes. This interval closely overlaps with that where we report topographical differences between groups. Importantly, our analysis provides new insights by showing that *different* neural mechanisms are involved when autistic and dyslexic children complete motion coherence tasks, compared to typically developing children: it is not that they just generally have increased or reduced activity in the same regions as typically developing children. Furthermore, our results suggest that increased activity over occipital electrodes (Toffoli et al., 2021) can be accompanied by overall reduced brain response strength in autistic and dyslexic children.

The second task that we investigated was an integration task, whereby, unlike the coherence task, there were no randomly moving noise dots. Instead, the dot directions in each stimulus were taken from a Gaussian distribution with higher (difficult) or lower (easy) variance. In contrast to the coherence task, the groups responded differently to the difficulty manipulation in this task. We first discuss the results for the two main effects. As in the coherence task, difficulty elicited early-onset topographic differences (110ms post-stimulus) that continued until the end of the epoch, and these were accompanied by mid-latency strength-based modulations (between ∼330-640ms), with stronger brain responses in the easy condition. However, in contrast to the coherence task, the early topographic modulations were also accompanied by early strength-based modulations (∼115-224ms post-stimulus), again stronger in the easy condition. At the group level, late strength-based effects were again found, albeit somewhat earlier (∼560-700ms vs. ∼630-800ms). In this task, significant brain response attenuations were found in both neurodivergent groups.

The group by task difficulty interaction in the integration task stemmed from differences in topographic mechanisms, at two separate time-periods. In the earlier time-period (434-500 ms), in the difficult condition, both autistic and dyslexic children differed from TD children, but in the easy condition, only the dyslexic children differed from TD children. In the later time-period (538-638ms), however, autistic children differed significantly from typically developing children in both the difficult and easy conditions, whereas dyslexic children showed only a strong trend towards a difference from TD children in the difficult condition (p = 0.05), and no difference in the easy condition. Despite these interactions, post-hoc comparisons showed no significant differences between autistic and dyslexic children, suggesting that these groups are more similar to each other in topographies than to typically developing children. Previous analyses based on univariate analyses of component waveforms (Toffoli et al., 2021) showed no group differences or interactions between group and difficulty in EEG activity in this task, which suggests that the topographical approach is more sensitive in identifying differences between typically developing and neurodivergent populations. Similarly, higher sensitivity of topographic analyses versus univariate analyses has been demonstrated in revealing age-based group differences in typically developing children (e.g., Turoman et al. 2021b,c) and differences within clinical pediatric populations as a function of treatment (e.g. Matusz et al. 2018).

Overall, our results show that autistic and dyslexic children’s brains respond differently to motion processing tasks than TD children. As in previous analyses (Toffoli et al., 2021), we report that group differences emerge only at relatively late processing stages (from ∼460 ms), with no evidence of group differences in early evoked responses to motion. Interestingly, dyslexic and autistic children followed a similar pattern of divergence from typical development in the coherence task, but showed subtly different patterns of divergence from typical development in the integration task (although there were no significant differences between autistic and dyslexic children). While both tasks involve discriminating the direction of global motion, their processing demands differ due to the distribution of dot directions: in the coherence task, there are randomly moving noise dots to be filtered out, whereas in the integration task, the optimal strategy is to average across all dot directions. Accordingly, our exploratory three-way analysis suggests, for the first time, that these tasks recruit distinct brain networks. Our results add to previous work showing that presenting these tasks in conjunction can be informative. Toffoli et al. (2021) reported group differences in a component maximal over occipital electrodes specifically for the coherence task (not the integration task) at mid-range latencies (∼430 ms), and speculated that this might reflect difficulties segregating signal-from-noise in both autism and dyslexia (e.g., Sperling et al., 2005; Zaidel et al., 2015). Our results here show that both autistic and dyslexic children use different brain mechanisms than typically developing children in the motion coherence task at similar mid-range latencies, which could potentially reflect different neural pathways relating to altered signal-from-noise segregation (e.g., atypical feedback connections; Raudies & Neumann, 2010).

Different patterns of diverging brain mechanisms in the direction integration task in autism and dyslexia could link to previously reported behaviourally-derived latent parameters in this task: increased sampling in autism (Manning et al., 2015; 2017) and increased internal noise in dyslexia (Manning, Hulks et al., 2022). However, this possibility would require testing in a study which incorporates both EEG and equivalent noise modelling. More generally, further work is needed to understand the relationships between differences in topographies and behaviour. Our behavioural data shows that dyslexic children were behaviourally less efficient than both autistic and dyslexic children in both tasks. This finding echoes that of reduced drift-rate in dyslexia (Manning et al., 2022b) but not autism (Manning et al., 2022a), which is not surprising, given that drift-rate, like inverse efficiency scores, combines both accuracy and response time. Given these behavioural differences, it is perhaps surprising that the differences in EEG between autistic and dyslexic children were non-significant. While the pattern of results vis-a-vis the typically developing children in the direction integration task suggests a potential, interesting difference between the autistic and dyslexic children in the motion integration task, the lack of direct differences between the groups necessitates further research.

Our results speak to an interesting possibility that there are both shared and non-shared effects on motion processing between autism and dyslexia (Manning, 2024), with both similar patterns, and subtle differences. Future work with larger sample sizes could allow greater sensitivity to detect differences between these conditions. Shared effects could reflect an underlying dorsal vulnerability (Braddick et al., 2003), or overlap in neural mechanisms across neurodevelopmental conditions more broadly (Siugzdaite et al., 2020). Here, the potential time constraints of effects to earlier versus later periods of late ERP activity between ∼400 and 650ms post-stimulus between the two groups, if replicated, could perhaps help with testing this hypothesis. At the same time, different patterns of divergence from typical development could suggest the need to work towards more condition-specific theoretical accounts. Extending this type of investigation to other types of neurodivergence (e.g., ADHD, Williams syndrome), will further help us understand how widespread topographical differences are. We note that our results do not provide support for one condition-specific account, namely, the magnocellular account of dyslexia (Stein, 2001), as group differences emerge at relatively late processing stages (although these tasks are suboptimal for assessing magnocellular functioning; Skottun, 2011).

While we have demonstrated differences in brain pathways in both autism and dyslexia, complementing our approach with techniques with increased spatial resolution will help better identify what precisely these brain pathways are. Previous fMRI studies have identified differences during coherent motion processing in autism and dyslexia, relative to typical development. In dyslexia, MT/V5 activation is reduced (Demb et al., 1998; Eden et al., 1996; Eden et al., 2003; Olulade et al., 2012). Meanwhile, in autism, increased activation in MT/V5 has been reported (Robertson et al., 2014; Takarae et al., 2014; Peiker et al., 2015). Based on these previous findings, we had expected that GFP might be increased in autism but reduced in dyslexia. However, we found no significant differences in autistic and dyslexic children’s GFP, which likely reflects that GFP is a marker of overall brain response strength, and not restricted just to MT/V5 (and indeed, increased activation in MT/V5 is not always reported [Koldewyn et al., 2011], and differences in V1 activation have also been reported [Brieber et al., 2010; Robertson et al., 2014]). These findings regarding specific regions of interest do not speak to differences in topographical organisation. Differences in functional connectivity have been reported in both autism (Müller et al., 2011) and dyslexia (Wang et al., 2026), which could relate to topographical differences, but again, we know of no work that has directly compared functional connectivity across the two conditions. Spiteri and Crewther (2021) recently proposed that motion processing differences in autism arise from atypical connectivity to the MT across development, whereby the typical developmental attenuation of the pathway from the pulvinar to MT/V5 could be slowed in autism. The authors also speculated that a similar account could explain motion processing in dyslexia, although these accounts require empirical testing.

The current study did not analyse differences in the recruited brain sources across the three groups, as the aim here was to identify topographic differences, and thus differences in the engaged brain regions, in the event-related EEG responses. The interesting potential differences in the temporality of the topographic mechanisms underlying how the autistic and dyslexic children process motion integration in the difficult conditions would need to be replicated and scrutinised with paradigms optimised for eliciting differences. Notably, electrical neuroimaging has been used to identify brain sources of differences between typically developing and neurodivergent populations (e.g. error or oddball processing in ADHD; Janssen et al. 2016, 2020). In subsequent studies optimised for identifying differences across the three populations, source estimations would be helpful in identifying the brain sources giving rise to the differences observed at the scalp level.

Longitudinal studies will shed further light into how motion processing develops in different developmental conditions. Hardiansiyah et al. (2023) recently reported that 5 month-olds with increased likelihood of autism who go on to have high levels of autistic characteristics at 36 months show different topographies in their EEG responses to coherent motion, with more lateralised processing across the visual cortex (stronger left lateralisation, and weaker at the midline) compared to infants with low likelihood of autism. This suggests that neurodiversity-related topographical differences may emerge very early in development. It would be informative to track these topographical differences from infancy to our mid-childhood age range, and to compare this trajectory with those who are at increased likelihood of, and who go on to develop, dyslexia.

## Conclusion

We report differences in the strength and topographies of autistic and dyslexic children’s brain activity during two motion processing tasks, compared to typically developing children, using an electrical neuroimaging approach. These findings extend previous findings from univariate approaches by showing that autistic and dyslexic children are not merely activating the same networks to a greater or lesser extent than typically developing children, but that they are employing different brain networks when completing motion tasks. Importantly, our electrical neuroimaging approach has helped to uncover neural differences in autistic and dyslexic children for a motion processing task that does not involve random, incoherent motion, demonstrating the sensitivity and utility of the approach for understanding neurodevelopmental diversity.

## Data availability statement

Anonymised data can be accessed through a safeguarded repository held by the UK Data Service: https://dx.doi.org/10.5255/UKDA-SN-855018. Users must register with the UK Data Service and sign the UK Data Service’s End User License Agreement. Experimental code can be found here: https://osf.io/wmtpx/.

## Ethics approval statement

The project was given ethical approval from the Central University Research Ethics Committee at University of Oxford.

## Participant consent statement

Parents of child participants gave written informed consent. Child participants gave written assent.

## Permission to reproduce material from other sources

N/A

## Supporting information

Supplemental figures S1-4 & their legends

## Acknowledgments

We are grateful to the participants and families who took part, the schools and organisations who kindly advertised the study, Irina Lepadatu, the Oxford Babylab and Dhea Bengardi for help with recruitment, and Lisa Toffoli, Helena Wood, Madeleine Mills, Amber Heaton, and Kate Seaborne who helped with data collection and data entry. The project was funded by a Sir Henry Wellcome Postdoctoral Fellowship awarded to CM (grant number 204685/Z/16/Z) and a James S. McDonnell Foundation Understanding Human Cognition Scholar Award to GS. CM is currently funded by the Medical Research Council (MR/Z504397/1). ES and PJM are funded by the Swiss National Science Foundation (Grant 10HW-9_220405). PJM is also funded by the Canton Valais (Grant PInter 13-2025) and Transforming Health & Care Systems 2024 (THCS no. 3230). AE was funded by a Rhodes Scholarship.

