## Supplemental figures S1-4 & their legends for "Topographical differences during motion processing in autistic and dyslexic children"

**Supplementary figures**

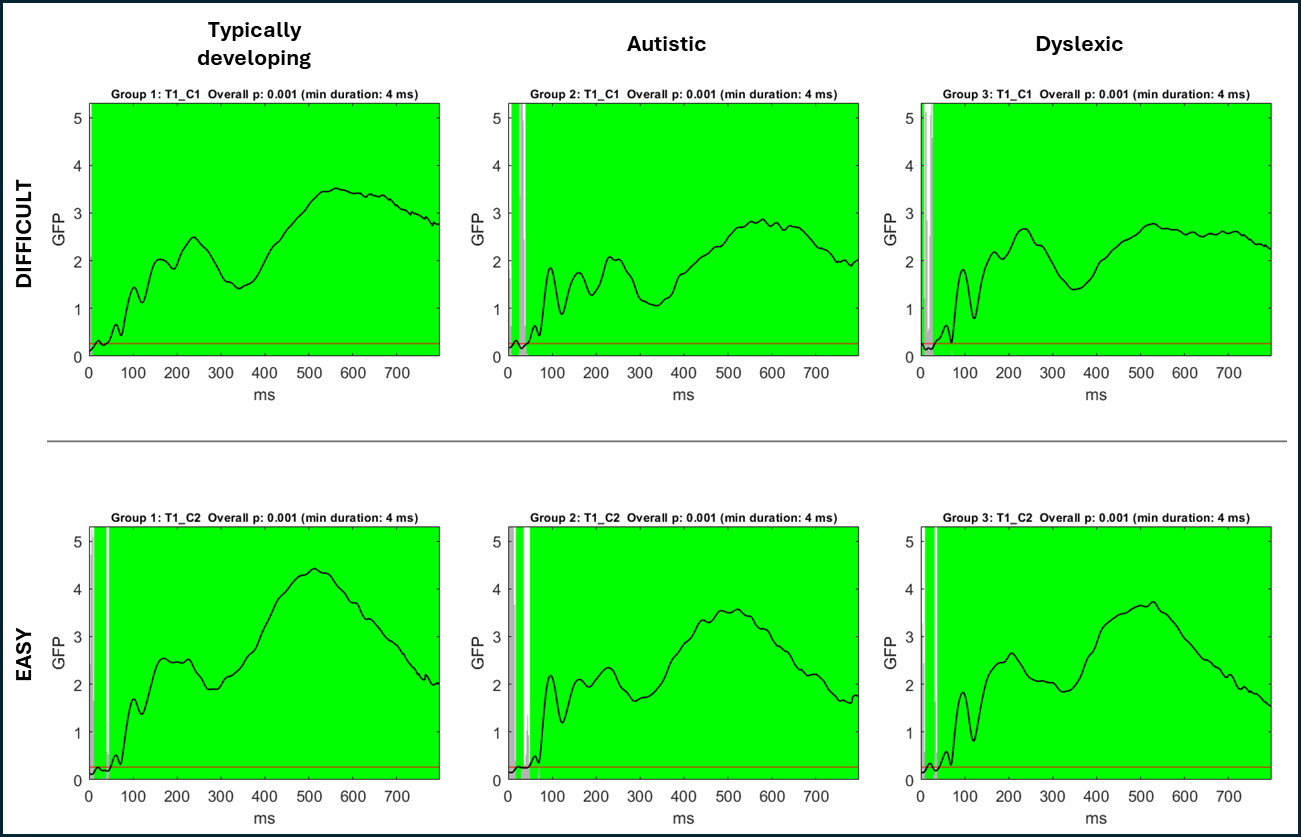

**Supplementary Figure 1**. TCT showed consistent neural activity in difficult (upper) and easy (bottom) conditions in the coherence task for all groups (typically developing, autistic and dyslexic).

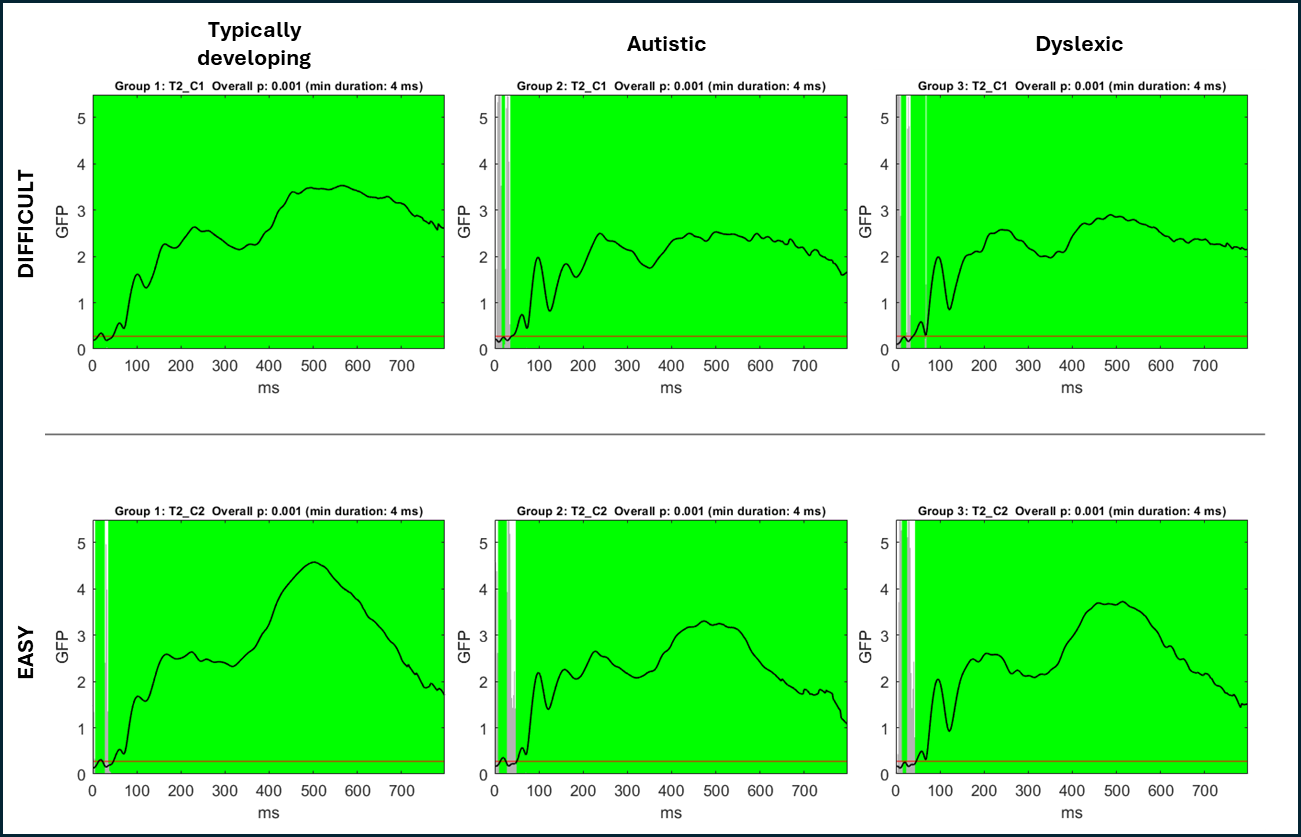

**Supplementary Figure 2.** TCT showed consistent neural activity in difficult (upper) and easy (bottom) conditions in the integration task for all groups (typically developing, autistic and dyslexic).

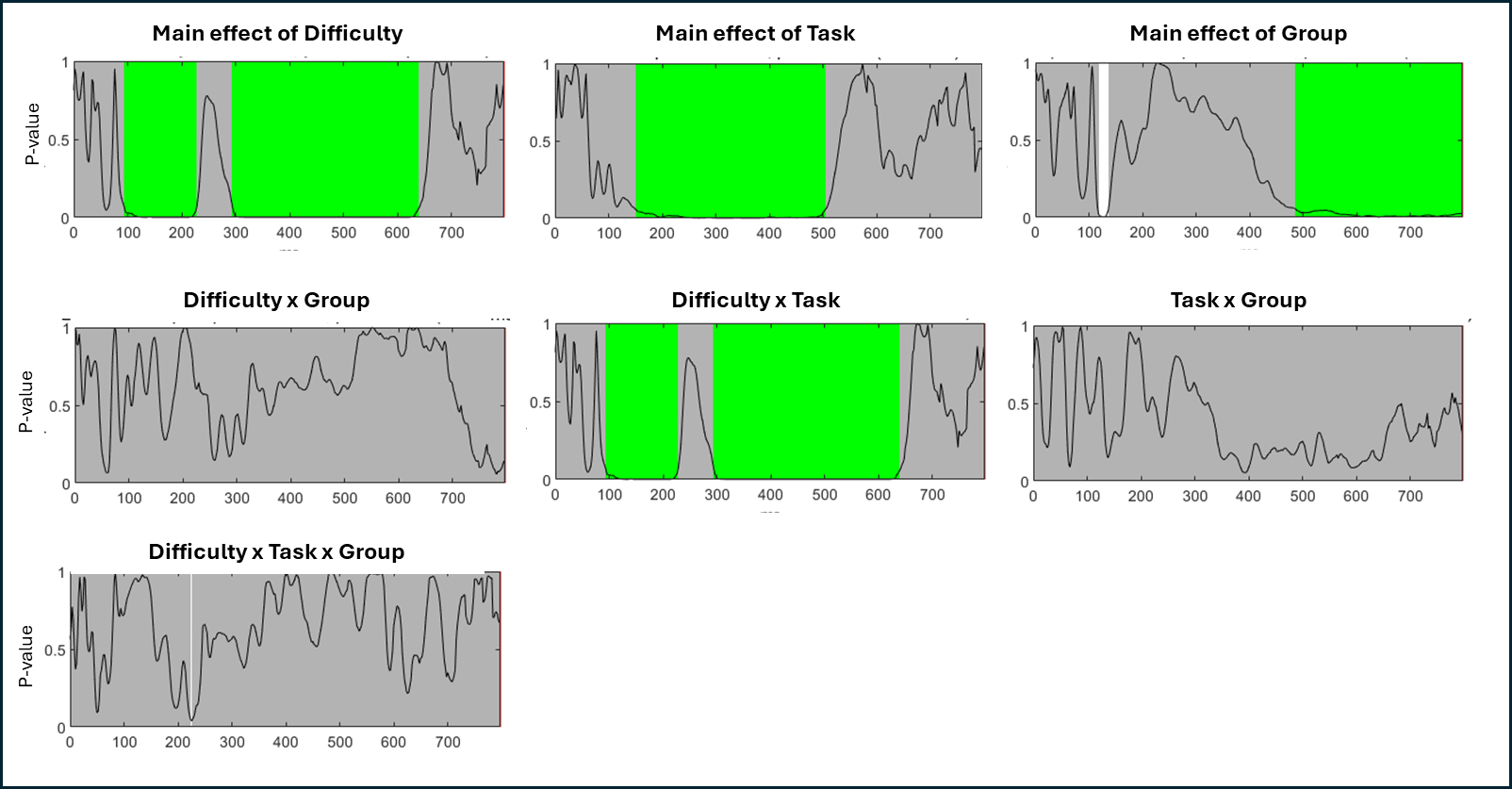

**Supplementary Figure 3.** GFP results of the three-way ANOVA are presented for the main effect of difficulty (difficult; easy), Task (coherence; integration) and group (typically developing; autistic; dyslexic) in whole-epoch analyses of the GFP. The black line represents the p-value, white areas indicate significant time points, and green segments mark intervals where the effect meets global duration criteria (i.e., GCS).

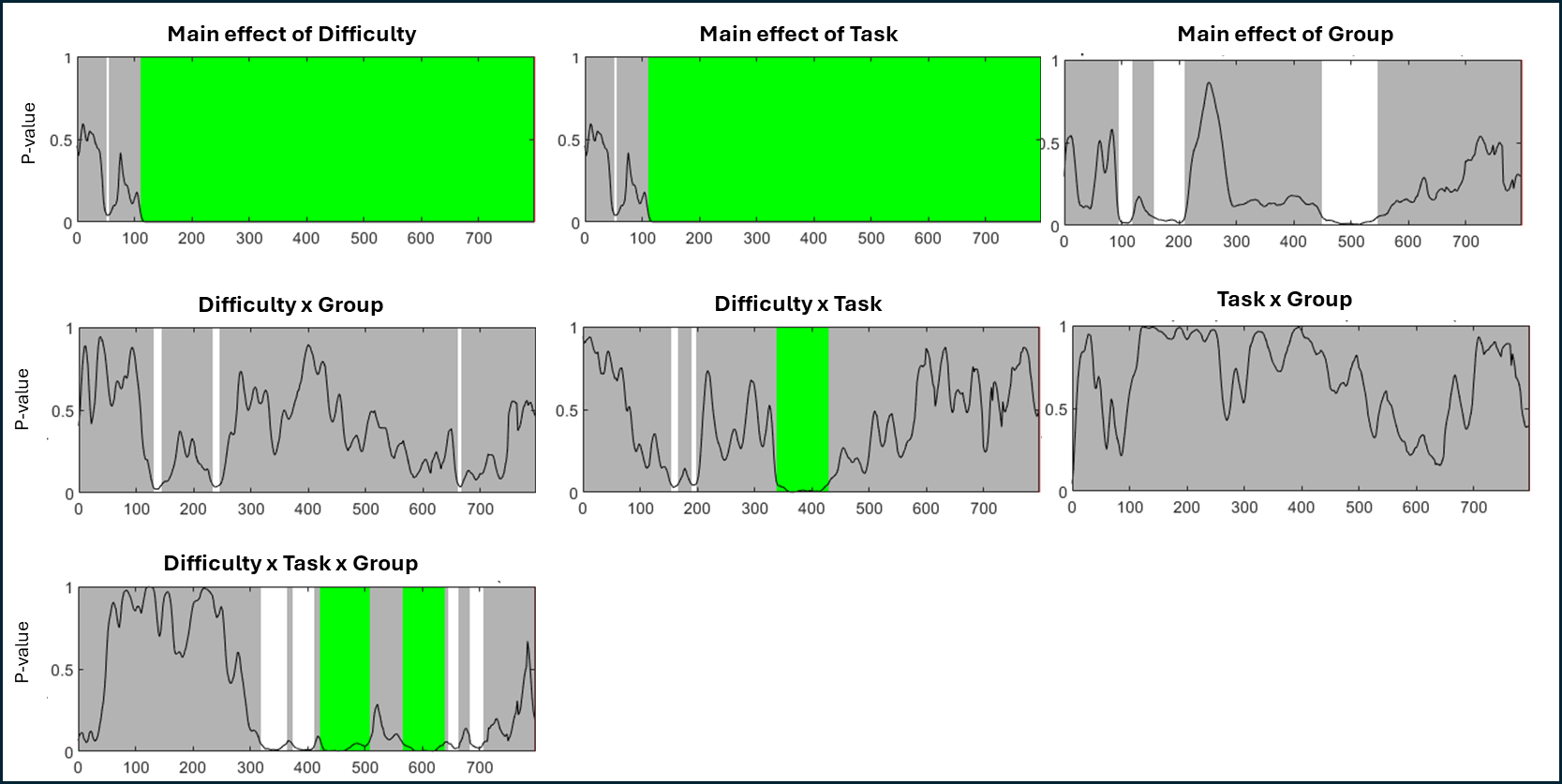

**Supplementary Figure 4.** Three-way TANOVA results**.** P-value plots are presented for the main effect of difficulty (difficult; easy), task (coherence; integration) and group (typically developing; autistic; dyslexic) in whole-epoch analyses of the TANOVA. The black line represents the p-value, white areas indicate significant time points, and green segments mark intervals where the effect meets global duration criteria (i.e., GCS).
